# Neural Mechanisms of Willed Attention Control

**DOI:** 10.64898/2025.12.22.696009

**Authors:** Changhao Xiong, Yujun Chen, Qiang Yang, Sungkean Kim, Sreenivasan Meyyappan, Jesse J. Bengson, George R Mangun, Mingzhou Ding

## Abstract

Cueing paradigms are commonly used to study the neural mechanisms of visual spatial attention control. In these paradigms, each trial starts with an external cue, which instructs the subject to pay covert attention to a spatial location in anticipation of an impending stimulus (instructed attention). Recent work has introduced a new type of cue which prompts the subject to spontaneously decide which spatial location to attend (willed attention). We studied the neural mechanisms of willed attention control by analyzing fMRI and EEG data recorded at two institutions (UF and UC Davis) using the same willed attention paradigm. The findings include: (1) both instructional cues and the choice cue activated the DAN, (2) the choice cue additionally activated a frontoparietal decision network consisting of dorsal anterior cingulate cortex (dACC), anterior insula (AI), anterior prefrontal cortex (APFC), dorsal lateral prefrontal cortex (DLPFC), and inferior parietal lobule (IPL), (3) the decision about where to attend can be decoded in frontoparietal decision network in choice trials but not in instructional trials, and (4) EEG alpha oscillation patterns immediately preceding the choice cue, but not the instructional cues, predicted the postcue direction of attention and the frontoparietal decision network activity. Based on these findings we proposed a model of willed attention control suggesting how the direction of visual spatial attention was decided upon in the absence of external instructions.

## Introduction

Visual spatial attention can be voluntarily directed in advance of stimulus processing to align with behavioral goals. The control of voluntary attention is often studied using cueing paradigms where each task trial begins with a cue that informs the participant where in visual space a task-relevant target stimulus may appear. This information enables participants to selectively prioritize stimuli at the cued location (Posner et al., 1980). The neural underpinnings of voluntary attention control have been extensively studied using cuing paradigms in humans using a multitude of techniques including patient lesion analyses (Posner et al., 1984), neuroimaging methods (Arrington et al., 2000; Corbetta et al., 2000; Hopfinger et al., 2000a; Woldorff et al., 2004) and electrophysiological and magnetoencephalographic recordings (Hopf & Mangun, 2000; Yang et al., 2024). Converging evidence shows that the cue activates the dorsal attention network (DAN), comprising the frontal eye fields (FEF) and the intraparietal sulcus (IPS), which then issues top-down control signals to bias sensory cortex to enhance relevant information and suppress irrelevant information (Corbetta & Shulman, 2002; Hopfinger et al., 2000b; Meyyappan et al., 2025), and the sensory biasing signals manifest as distinct patterns of alpha oscillations (Worden et al., 2000).

In order to study voluntary attention control in more realistic circumstances, another method has been introduced. In this method, instead of presenting cues that tell the subject where a relevant item is likely to appear or instructs them where to focus attention, subjects are permitted to freely choose where to attend (Taylor et al., 2008). These so-called “choice cues” prompt the subject to spontaneously decide where to attend among a set of alternatives (e.g., left visual field vs. right visual field locations). This form of voluntary attention has been termed “willed attention” (Bengson et al., 2015). Studies of willed attention control have shown that in addition to DAN, the choice cue elicits activation in an additional set of frontoparietal areas, including DLPFC, IPL, dACC and anterior insula (Bengson et al., 2015; Hopfinger et al., 2010; Rajan et al., 2019; Taylor et al., 2008). The functional role of these frontoparietal areas, however, has yet to be clarified. Given that prior work has shown that these areas are involved in decision-making, it is thus reasonable to hypothesize that, in willed attention, these areas may contribute to the internally generated decision about where to attend. The first goal of this study is to test whether activity patterns in this network are consistent with representing the content of that decision.

The control of willed attention begins with the decision about where to direct spatial attention when prompted by the choice cue, which appears unpredictably both in time and among the three cue types (attend-left, attend-right, choose). Previous work on the neuroscience of decision-making has shown that the ongoing brain state before the stimulus can exert a significant influence on the outcome of the decision-making process. A key variable indexing the ongoing brain state is the alpha oscillation (8-12 Hz) (Jensen & Mazaheri, 2010; Klimesch et al., 2007). In stimulus-based decision-making, prestimulus alpha power is shown to mainly affect the response criterion. When prestimulus alpha power is low, individuals exhibit a more liberal detection criterion, manifesting as an increased tendency to report stimulus presence even when the evidence is ambiguous (Limbach & Corballis, 2016). Conversely, higher pre-stimulus alpha power promotes a more conservative response strategy, characterized by increased “no” responses and higher decision thresholds (Iemi et al., 2017).

In typical willed attention studies, however, the decision is between two equal alternatives, attend-left versus attend-right visual field locations, and the choice cue offers no sensory information that favors either one. The previous study (Bengson et al., 2014) reported that the amplitude of left parieto-occipital EEG alpha power in the 1 sec period immediately before the unpredictable appearance of the choice cue predicted the decision about where to attend. How far does this alpha-based predictability extend before the choice cue? Bengson and colleagues (2014) investigated multiple sequential 1 sec periods prior to the choice cue and reported no significant predictive value of parieto-occipital alpha power in earlier time windows. But this analysis had rather low temporal resolution, relying on alpha power across rather large and arbitrary time windows. Knowing more about the timecourse of the relation between pre-choice-cue alpha power and behavioral choices would help assess whether predictive information is concentrated near cue onset, consistent with momentary brain-state biases, or instead extends over a longer interval, consistent with longer-lasting biases in brain activity patterns. In addition, understanding how precue alpha-power patterns relate to subsequent choice-cue-evoked neural activity in the frontoparietal decision network would offer deeper insight into the role of the ongoing brain state in shaping decision-making during willed attention. The second goal of this study, therefore, is to address these questions.

Healthy volunteers performed a cued visual spatial willed attention paradigm, in which each trial started with one of three types of cues: two instructing the participant to covertly attend the left or the right visual field and the third prompting the participant to spontaneously choose between the left and the right visual field to covertly attend. Following a random time delay, a stimulus was presented, and the participant discriminated features of the stimulus if it appeared in the attended visual field and ignored it completely if it appeared in the unattended visual field. EEG and fMRI data were recorded at two different sites, University of Florida (UF) and University of California at Davis (UCD), using the same paradigm. Data from the two sites were analyzed separately to assess the consistency of the principal findings and then combined at a higher-level analysis to increase statistical power.

## Materials and Methods

### Overview

Two independent datasets were recorded at two different sites, University of Florida (UF) and University of California Davis (UCD), using the same experimental paradigm. These datasets have been analyzed and published on before to address different questions (Bengson et al., 2015; Liu et al., 2017; Meyyappan et al., 2025). None of the previously published studies examined the questions considered here.

#### UF Dataset

The experimental protocol was approved by the Institutional Review Board of the University of Florida. A total of 18 right-handed college students whose age ranged between 18-22 gave written informed consent and participated in the study. They had normal or corrected to normal vision and no history of neurological or psychological disorders. Data from five participants were excluded from the analysis based on the following reasons: (1) the behavioral performance was below criterion (< 70% accuracy, one participant); (2) unable to follow the task instructions (one participant); and (3) excessive body or eye movements in the scanner (three participants). The remaining thirteen participants analyzed here included 5 females and 8 males.

#### UCD Dataset

The experimental protocol was approved by the Institutional Review Board of the University of California, Davis. Overall, 19 right-handed college students whose age ranged between 18-22 gave written informed consent and participated in the study. All participants had normal or corrected to normal vision and had no history of neurological or psychological disorders. One participant was excluded due to unstable performance and failure to follow instructions during the experiment. The remaining eighteen participants were analyzed which included 2 females and 16 males. The sex distribution in the UCD sample was imbalanced; the sample size and imbalance did not permit a meaningful examination of sex-related effects.

### Paradigm and procedure

See Figure 1. Each trial began with one of three symbolic cues (circle, square, or triangle) displayed slightly above the central fixation for a duration of 200 ms. Two of the cues, called instructional cues, required the participant to covertly direct their attention to the left or right visual field while maintaining central fixation (instructed attention). The third cue, called the choice cue, prompted the participant to freely choose one of the two visual fields to attend while maintaining central fixation (willed attention). The three cue conditions occurred with equal probability and the symbols used as cues were counterbalanced as to their meanings across participants. Further, the willed and the instructed attention trials were randomly interleaved in order to discourage stereotypic responses. Following a cue-target interval randomized between 2000 and 8000 ms, a black and white grating of two possible spatial frequencies (low vs. high) appeared in one of the two visual fields with equal probability for 100 ms (50% target validity). Participants discriminated the spatial frequency of the target grating appearing in the attended visual field as fast and accurately as possible through a button-press and ignored the grating appearing in the unattended visual field. The gratings were difficult to discriminate so that focused spatial attention was required for successful execution of the task. Compared to the probabilistic cueing used in the Posner style paradigms (Posner, 1980), in which participants responded to the stimulus appearing in either visual field, the current design further encouraged the full deployment of focused covert spatial attention prior to the appearance of the stimulus.

**Figure 1.**
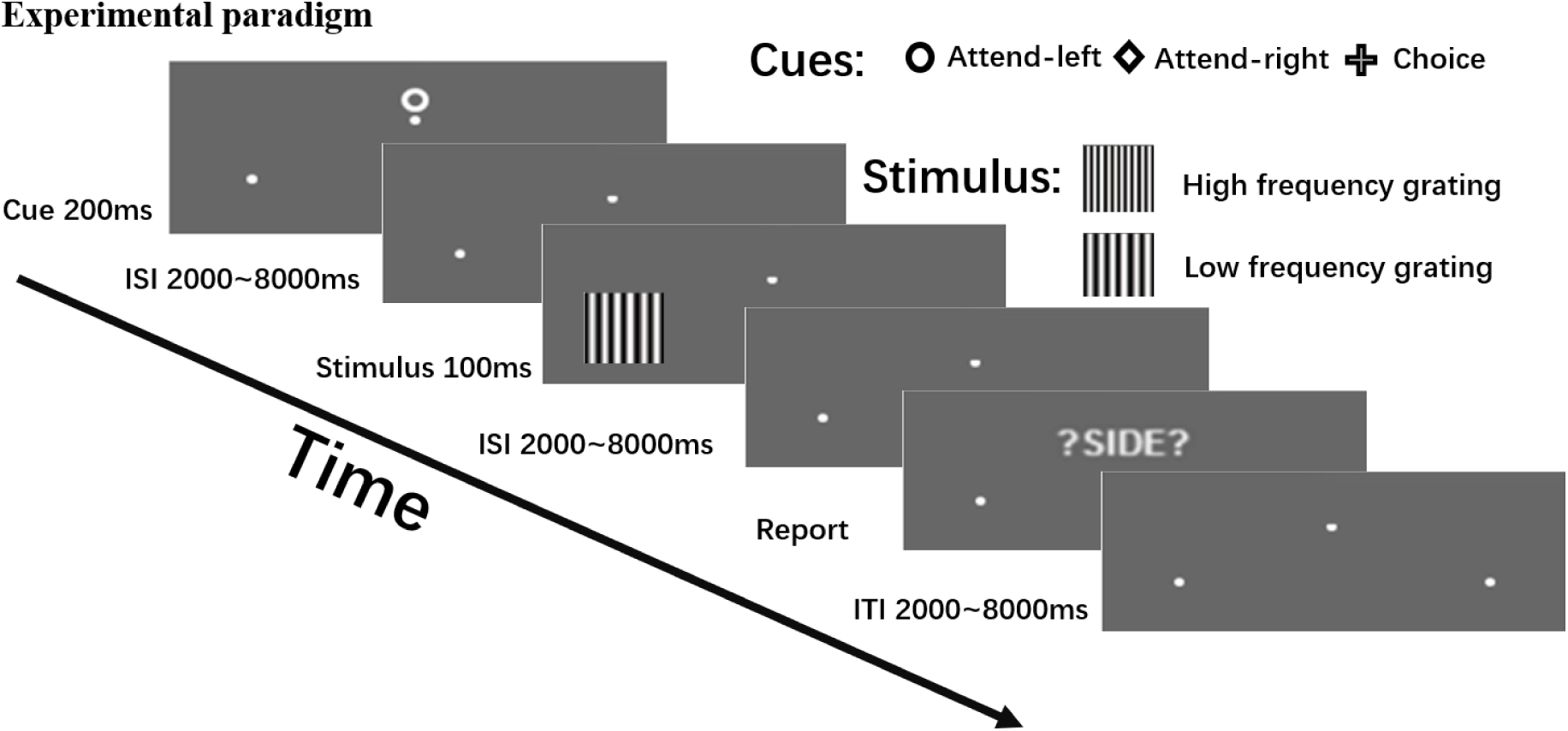
Experimental paradigm. Each trial began with the presentation of one of three visual cues for 200 ms. Two of the cues—the instructional cues— instructed the participant to covertly attend to either the left or right visual field, the third cue—the choice cue—prompted participants to choose whether to attend to the left or right field. Shown in the figure is an example of the cue symbol-to-attention mappings which were randomized across participants. After a variable cue-to-target interval (2,000–8,000 ms), a target stimulus (a grating with high or low spatial frequency) appeared briefly (100 ms) with equal probability in either visual field. Participants were asked to discriminate the spatial frequency of the grating presented at the cued or chosen location (attended location), while ignoring stimuli on the uncued or unchosen side (unattended location). Following a variable inter-stimulus interval (ISI; 2,000–8,000 ms), a “?SIDE?” prompt appeared, asking participants to report the visual field they attended to on that trial. Finally, a variable inter-trial interval (ITI; 2,000–8,000 ms) elapsed before the onset of the next trial.

Following an inter-stimulus interval (ISI) randomized between 2000 and 8000 ms after the stimulus onset, participants were prompted by the visual cue “?SIDE?” to report the visual field they were attending on that trial through a button-press (self-report). This reporting was used to determine the direction of attention for willed attention trials but was included in instructed attention trials as well to maintain trial structure consistency. Reporting of the attended side was highly accurate across the participants with the accuracy of 98.4% averaged across conditions. For instructed attention trials, side reporting accuracy could be determined by comparing the self-report with the cue instruction, while for willed attention trials, the accuracy was determined by examining the consistency between the participant’s response (or lack of response) and the self-report. An inter-trial interval (ITI) randomized between 2000 and 8000 ms was inserted after the self-report. This variable ITI generated uncertainty about when the next cue would occur and was intended to discourage participants from adopting a fixed advance-selection strategy.

Prior to recording, all participants went through a training session to familiarize with the task and to ensure stable performance (above 70% accuracy). During recording, the experiment was divided into blocks of trials with the length of each block kept around six minutes. All participants completed at least eight blocks of trials. For willed attention, participants were explicitly told to avoid relying on any stereotypical strategies of generating decisions, such as always attending the same/opposite side they attended during the previous trial, as well as to avoid randomizing or equalizing their decisions to choose left or right across trials; prior studies found that decisions to explicitly randomize decisions might invoke additional working memory related processes (Spence & Frith, 1999).

### Data collection

#### UF fMRI

fMRI data were acquired using a 3T Philips Achieva scanner (Philips Medical Systems, Netherlands) with a 32-channel head coil. Functional images were obtained using a single-shot echo planar imaging (EPI) sequence with the following parameters: repetition time (TR) = 1980 ms, echo time (TE) = 30 ms, flip angle = 80°, field of view = 224 mm. Thirty-six transverse slices (slice thickness: 3.5 mm) oriented parallel to the plane connecting the anterior and posterior commissures were obtained in an ascending order with a voxel resolution of 3.5×3.5×3.5 mm^3^.

#### UF EEG

Continuous EEG data was recorded inside the MRI scanner simultaneously with fMRI using a 32-channel MR-compatible EEG recording system (Brain Products, Germany). Thirty-one Ag/AgCl electrodes were located on the scalp according to the 10–20 system via an elastic cap. One additional electrode was located on the participant’s upper back to record the electrocardiogram (ECG), which was used to remove the ballistocaridogram (BCG) artifacts (Ertl et al. 2010). The FCz channel was used as a reference during recording. The impedance of all scalp channels was kept under 5 kΩ throughout the experiment. The sampling frequency was 5000 Hz.

#### UCD fMRI

fMRI data were acquired using a 3T Siemens Skyra scanner (Siemens, Germany) with a 32-channel head coil. Functional images were obtained using a single-shot EPI sequence with the following parameters: TR = 2100 ms, TE = 30 ms, flip angle = 90o, field of view = 218 mm, slice thickness = 3.4 mm. Thirty-four transverse slices were obtained in an interleaved acquisition order. The voxel resolution was 3.5×3.5×3.5 mm^3^.

#### UCD EEG

Continuous EEG data was recorded in a separate session from 64 electrodes mounted in an elastic cap (Electro-cap International Inc., OH) using a SynAmps II amplifier (Compumedics Neuroscan, USA). The reference electrode was placed at the right mastoid during recording. The impedance of all scalp channels was kept below 5 kΩ. The sampling frequency was 1000 Hz.

Here, site-specific acquisition factors, including scanner parameters, EEG montage, and simultaneous versus separate EEG/fMRI acquisition, were not modeled as moderators. Primary analyses were conducted within subjects and separately by dataset before higher-level combination.

### fMRI data preprocessing

The preprocessing of fMRI data was performed with SPM8 and included the following steps: slice time correction, motion realignment, spatial normalization and smoothing. Slice timing correction was carried out using sinc interpolation. Motion realignment was performed by co-registering all images with the first scan of each session. Six motion parameters (3 translational and 3 rotational) were calculated and entered into the final design matrix to be regressed out in the general linear model (GLM) analysis. All images were spatially normalized to the standard MNI space and resampled to a voxel resolution of 3×3×3 mm. Finally, the spatially normalized images were smoothed using a Gaussian kernel of 8 mm full width at half maximum. A high pass filter with a cut off frequency of 1/128 Hz was used to remove any low frequency noise. Global effects were adjusted by the proportional scaling approach (Fox et al., 2009).

### EEG data preprocessing

For the UF dataset, since EEG data were recorded simultaneously with fMRI inside the scanner, extra preprocessing steps were required to remove the MRI artifacts. Both gradient and ballistocardiogram (BCG) artifacts were corrected via a template based average artifact subtraction method (Allen et al. 1998, 2000) implemented in Brain Vision Analyzer 2.0 (Brain Products, Germany). The gradient artifacts were corrected by constructing an average artifact template over 41 consecutive volumes in a sliding-window fashion and then subtracting this template from the raw EEG data for each volume. For the BCG artifacts, ECG R-waves were first detected, and 21 consecutive ECG segments defined around the R-waves were averaged to produce a BCG artifact template. This template was then subtracted from the EEG data to correct for BCG contamination.

After removing the MRI artifacts, the preprocessing and data analysis procedures for UF and UCD EEG datasets were the same, both using EEGLAB. EEG data were down sampled to 250 Hz and bandpass filtered between 0.1 and 50 Hz. Two types of epoching were carried out. In the first type, the bandpass filtered continuous EEG data were epoched from 500 ms before cue onset to 1500 ms after the cue onset, namely, −500 ms to 1500 ms. In the second type, the bandpass filtered continuous EEG data were epoched from −2500 ms before the cue onset to the cue onset, namely, −2500 ms to 0 ms. The data epoched from −500 ms to 1500 ms were used to examine neural activity immediately preceding the cue onset, the cue-evoked responses, and how the two might related to one another, whereas the data epoched from −2500 ms to 0 ms were used to examine whether neural activity farther ahead of the cue onset could impact the decisions and cue-evoked responses. The epoched data were visually inspected and trials with large artifacts were rejected. Independent component analysis was then applied to remove EEG components related to eye blinks, eye movements and other artifacts. It is worth noting that since the intertrial intervals (ITIs) varied between 2000 ms to 8000 ms, trials whose cue onset occurred within 3000 ms of the end of the previous trial were not considered for the second type of epoching, resulting in fewer trials (UF: choose-left = 36 ± 11, choose-right = 34 ± 10, instructed-left = 64 ± 17, instructed-right = 64 ± 15; UCD: choose-left = 46 ± 13, choose-right = 52 ± 16, instructed-left = 92 ± 17, instructed-right = 91 ± 19).

### fMRI activation analysis

Univariate BOLD activation analysis was carried out using the general linear model (GLM) method. Seven regressors were used to model the relevant events during the experiment: two for instructional cues (cue-left and cue-right), two for choice-left and choice-right, one for incorrect trials, and two for the stimulus appearing in the left and the right visual field. The cue-evoked activation was obtained for each voxel within each subject by applying t-test to the appropriate regressors. The group-level activation map was obtained by performing a one-sample t-test on each subject’s contrast maps with a threshold of p < 0.05 correcting for multiple comparisons with false discovery rate (FDR).

### fMRI multivariate pattern analysis

Multivariate pattern analysis (MVPA) was performed at the level of region of interest (ROI) using the linear support vector machine (SVM) method implemented in the LIBSVM package (http://www.csie.ntu.edu.tw/~cjlin/libsvm/) (Chang & Lin, 2011) with c = 1. ROIs were defined as 5-mm-radius spheres centered on peak coordinates from the group-level choice > instructed contrast (Table 2). This selection contrast collapsed across left and right trials within each condition, whereas MVPA decoded left versus right trials. Thus, the ROI-selection contrast was not based on the left-right distinction tested in the decoding analysis. For decoding analyses beyond the ROIs considered here, see Meyyappan et al. (2025), which showed that left-versus-right attention can be decoded in the DAN and throughout the visual hierarchy in both choice and instructed trials.

The trial-by-trial BOLD estimate following cue onset was obtained using the beta series regression method (Rissman et al., 2004). This method involved modeling each trial with a separate regressor in the GLM and the beta coefficients represented the estimated single-trial BOLD responses to the cue. Thus, the fMRI MVPA was performed on one cue-locked single-trial beta estimate per trial rather than on individual TRs or a time-resolved fMRI signal. The beta coefficient of each voxel in a participant was z-normalized across trials to remove any intrinsic trend. For decoding, cue-related data were normalized within each ROI by subtracting the voxel-wise mean and dividing by the voxel-wise standard deviation, which removed the mean activation level and allowed the focus on pattern information. A 10-fold cross-validation procedure was then employed where the cross-validation folds were randomized across the entire session rather than blocked by run; it is worth noting that this procedure may not fully eliminate run-specific autocorrelation or scanner-drift effects. A SVM classifier was trained on nine folds of the data and tested on the remaining fold. This process was repeated 10 times, each time using a different subset of trials for testing, and the decoding accuracies were then averaged across folds. To improve the robustness, the entire 10-fold cross-validation procedure was repeated 100 times with different random fold assignments of the data. The decoding accuracy for each ROI was computed by averaging across all repetitions. This decoding accuracy was then averaged across participants to obtain the group-level decoding accuracy. Decoding accuracy was considered significant if it exceeded 50% chance level using a group-level one-sample t-test across participants; FDR correction was applied to control for multiple comparisons across ROIs. Above-chance decoding accuracy was interpreted as evidence for the presence of task-relevant information in the ROI.

### EEG multivariate pattern analysis

One of the main foci of this study was on whether multi-electrode patterns of alpha power in the time period prior to cue onset predicted the postcue decision about where to attend, and what the time course of those predictive patterns might be. For the EEG data epoched from −500 ms to 1500 ms, with 0 ms denoting cue onset, trial-by-trial alpha power (8-12 Hz) were calculated over −500 ms to 0 ms interval using the Fast Fourier transformation (FFT). The power of other frequencies between 5 Hz and 25 Hz was also calculated over −500 ms to 0 ms interval. For the EEG data epoched from −2500 ms to 0 ms, we computed the alpha power as a time series from −2500 ms to 0 ms using a moving-window approach, where the window duration was 500 ms and the step size was 20 ms. The time of the window was labeled by the time of the rightmost point (e.g., the time of the window −2500 ms to −2000 ms was labeled as 2000 ms).

The MVPA on the EEG data was also performed using linear support vector machine (SVM) method implemented in the LibSVM package (http://www.csie.ntu.edu.tw/~cjlin/libsvm/) (Chang & Lin, 2011) with c = 1. The pattern of alpha power across electrodes was used as the input feature to predict the attended spatial location (left vs. right) after the cue. The classification accuracy for each participant was calculated using a 10-fold cross validation technique where the cross-validation folds were randomized across the entire session rather than blocked by run. Nine folds of the labeled data (i.e., attend-left and attend-right; choose-left and choose-right) were randomly chosen and used for training a classifier to generate a predictive model. The remaining one fold of the data was used to test the model by comparing the actual labels against the predicted labels. This process was repeated 100 times and the prediction accuracy from each 10-fold partition was averaged to yield the final decoding accuracy. For the EEG data epoched from −500 ms to 1500 ms, this analysis produced a single decoding accuracy, which informed whether precue alpha power pattern in (−500 ms to 0 ms) can predict the postcue direction of attention. Decoding accuracy was considered significant if it exceeded 50% chance level using a group-level one-sample t-test across participants. For the EEG data epoched from −2500 ms to 0 ms, the moving-window decoding analysis produced a time series of decoding accuracy, and a time-point-by-time-point statistical test was carried out to determine at which time point prior to cue onset, the pattern of alpha activity began to predict the postcue direction of attention. To further control for family-wise error across the time course, a cluster-based random permutation procedure with 10,000 permutations was applied.

### Relating precue EEG activity with postcue fMRI activity

To examine the relation between precue alpha power patterns and cue-evoked fMRI activity, we divided the participants into two groups via a median split in terms of precue alpha decoding accuracy: *low* precue alpha decoding accuracy group and *high* precue alpha decoding group. We then compared cue-evoked BOLD activation and BOLD decoding accuracy in the frontoparietal decision network between the two groups, including ROIs in the dACC, aPFC, DLPFC, IPL, and AI. As an exploratory, descriptive summary measure, we calculated a neural-efficiency index combining BOLD activation and decoding accuracy:

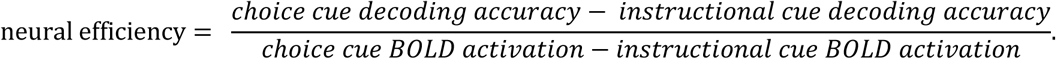

In this definition, univariate BOLD activation in the denominator is used to reflect the overall metabolic activity underlying the decision about where to attend, whereas the decoding accuracy in the numerator is used to indicate the strength of the decision about where to attend. Larger numerators coupled with smaller denominators yield higher neural efficiency, namely, participants who exhibit clearer decision-related frontoparietal brain patterns while consuming less energy in doing so are more efficient decision-makers. It is worth noting that whereas the notion of neural efficiency is not new (Dunst et al., 2014; Haier et al., 1988), the definition above, however, has not been attempted in the past. We also caution that, because ratio-based measures can be unstable when their denominator is small or noisy, this index was treated as a suggestive measure rather than the main outcome measure.

For statistical analysis, we conducted bootstrap resampling on both the BOLD signal differences and the decoding accuracy differences using 10,000 iterations, drawing, with replacement, samples equal in size to the original dataset and computing the mean for each resample. This yielded a bootstrap distribution consisting of 10,000 sample means. Higher values indicate greater choice-related decoding differences relative to choice-related BOLD-activation differences.

### Combining the two datasets via meta-analysis

The UF dataset and the UCD dataset were separately analyzed, and the results were examined for consistency and reproducibility. Where appropriate, p-values from the two datasets were subsequently combined using the Liptak-Stouffer method (Lipták, 1958). This method has been used to combine results from multiple datasets to enhance statistical power and rigor (Cheng et al., 2015; Huang & Ding, 2016; Liu et al., 2017). In this method, the following procedure was used to obtain the meta-analysis p values for the statistical tests. First, the p value of each dataset was transformed into Z-score according to the formula: *Z*= Φ^−1^(1−p), where Φ denotes the standard normal cumulative distribution function. Second, the Liptak-Stouffer *Z*-score of the meta-analysis level was calculated according to the formula (Lipták, 1958):

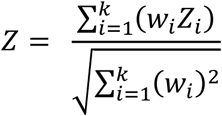

where *Z* is the meta-analysis *Z*-score of the two different datasets (here, *k*=2), *i* refers to the *i*th dataset, *w_i_* = √*N_i_* is the weight of the *i*-th dataset, and *N_i_* is the number of participants in *i*th dataset. Finally, the p values of the meta-analysis were obtained from the Liptak-Stouffer *Z*-scores. Here, meta-analysis was used to combine inferential results across datasets and did not model site-specific acquisition factors as moderators.

## Results

### Behavioral analysis

#### UF dataset

See Table 1. For cue-left and cue-right trials, the reaction times (RTs) to valid targets were 918.56 ± 24.49 ms and 911.81 ± 23.28 ms and the target discrimination accuracies were 85.72 ± 2.14% and 85.80 ± 1.24%, respectively. There were no statistically significant differences between cue-left and cue-right conditions in either RT (p = 0.67) or accuracy (p = 0.97). For choose-left and choose-right trials, RTs to valid targets were 942.27 ± 47.75 ms and 938.25 ± 29.04 ms, and accuracies were 88.41 ± 1.88% and 90.78 ± 1.28%, respectively. Statistical comparisons revealed no significant differences were found between conditions in either RT (p = 0.93) or accuracy (p = 0.25).

**Table 1.**
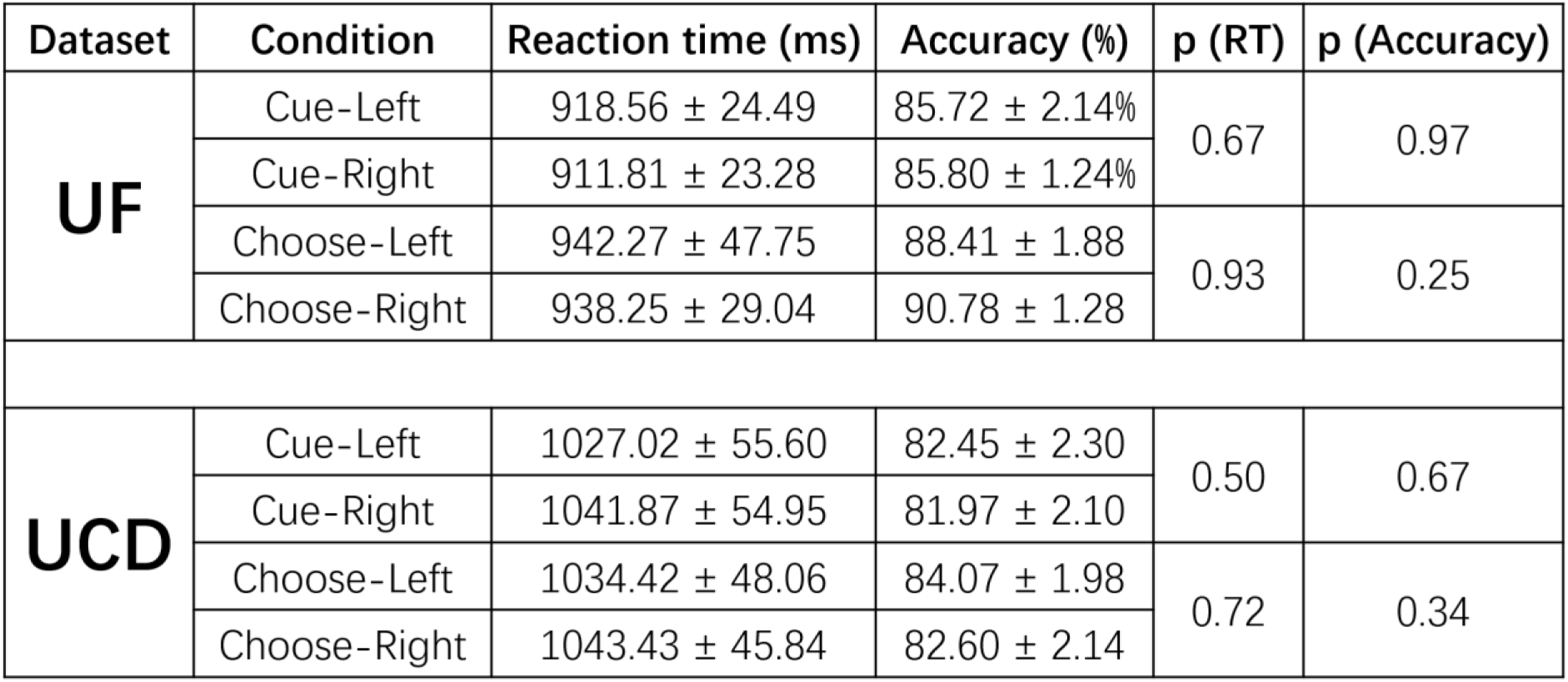
Behavioral results for UF and UCD datasets.

| Dataset | Condition | Reaction time (ms) | Accuracy (%) | p (RT) | p (Accuracy) |
| --- | --- | --- | --- | --- | --- |
| UF | Cue-Left | 918.56 ± 24.49 | 85.72 ± 2.14% | 0.67 | 0.97 |
|  | Cue-Right | 911.81 ± 23.28 | 85.80 ± 1.24% |  |  |
|  | Choose-Left | 942.27 ± 47.75 | 88.41 ± 1.88 | 0.93 | 0.25 |
|  | Choose-Right | 938.25 ± 29.04 | 90.78 ± 1.28 |  |  |
| UCD | Cue-Left | 1027.02 ± 55.60 | 82.45 ± 2.30 | 0.50 | 0.67 |
|  | Cue-Right | 1041.87 ± 54.95 | 81.97 ± 2.10 |  |  |
|  | Choose-Left | 1034.42 ± 48.06 | 84.07 ± 1.98 | 0.72 | 0.34 |
|  | Choose-Right | 1043.43 ± 45.84 | 82.60 ± 2.14 |  |  |

**Table 2.**
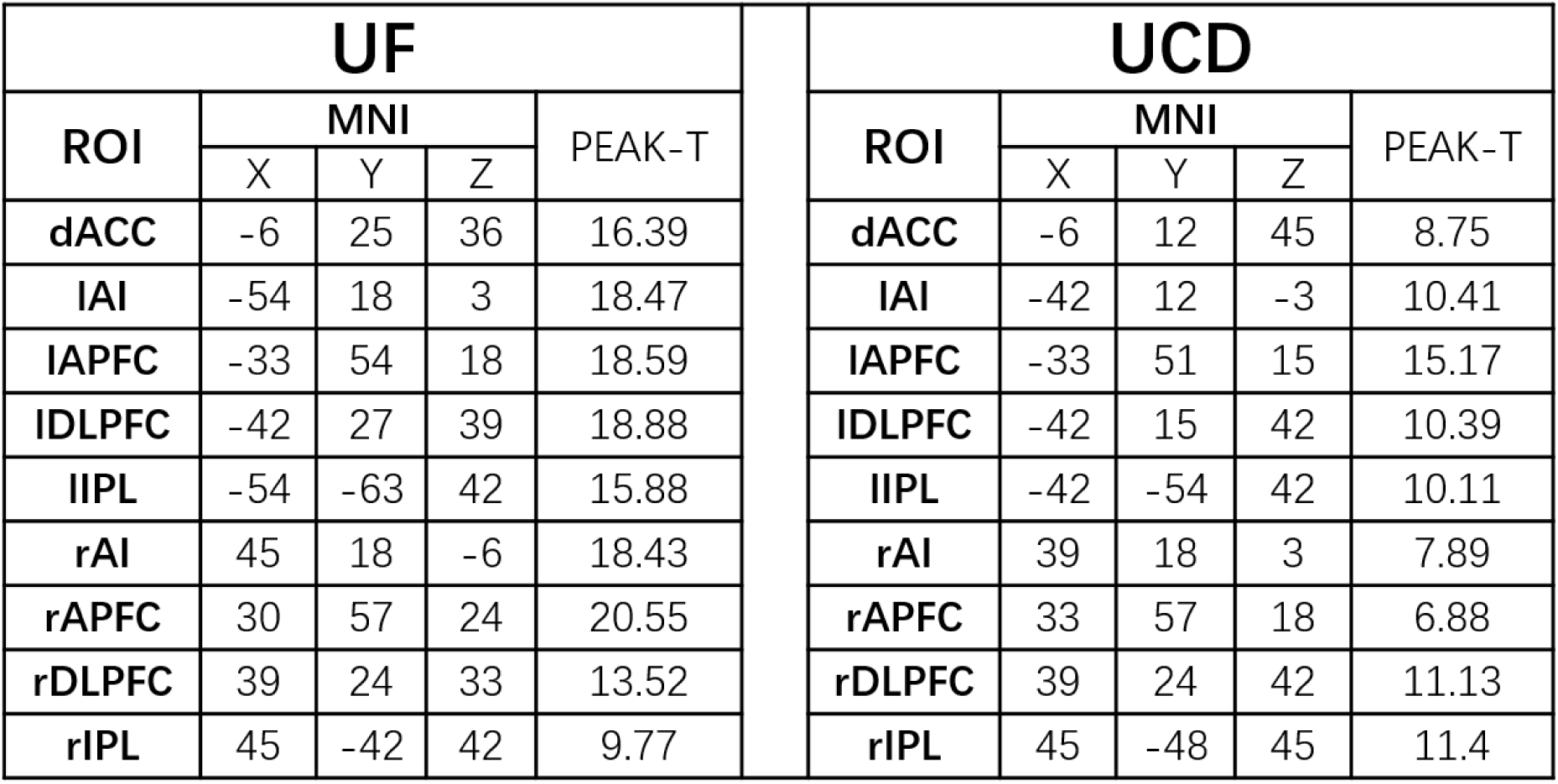
Frontoparietal regions preferentially activated by choice cues relative to instructional cues across both datasets. MNI coordinates of the contrast peaks are shown. Regions of interest (ROIs) were defined as 5-mm-radius spheres centered on these peaks.

#### UCD dataset

See Table 1. For cue-left and cue-right trials, RTs to valid targets were 1027.02 ± 55.60 ms and 1041.87 ± 54.95 ms and accuracies were 82.45 ± 2.30% and 81.97 ± 2.10%, respectively. No significant differences were observed between the two attention conditions for either RT (p = 0.50) or accuracy (p = 0.67). For choose-left and choose-right trials, RTs were 1034.42 ± 48.06 ms and 1043.43 ± 45.84 ms and accuracies were 84.07 ± 1.98% and 82.60 ± 2.14%, respectively. Statistical comparisons revealed no significant differences in RT (p = 0.72) or accuracy (p = 0.34).

It is worth noting that behavioral performance differed quantitatively between the UF and UCD cohorts, with UCD participants showing slower RTs and somewhat lower accuracy despite use of the same willed-attention paradigm and matched stimulus parameters. The source of these differences is unknown. One possible contributor is that EEG and fMRI were recorded simultaneously in a single session at UF but in separate sessions at UCD, which may have affected task engagement or fatigue. In addition, relative to matched instructed trials (choose-left versus cue-left; choose-right versus cue-right), choice trials showed numerically longer RTs and higher accuracy across the two datasets. However, these effects were not consistently significant across measures and sites (UF: RT, p = 0.438; accuracy, p = 0.012; UCD: RT, p = 0.830; accuracy, p = 0.292). This descriptive pattern may be consistent with a speed-accuracy trade-off associated with increased cognitive control.

### Univariate analysis of cue-evoked BOLD activation

Both the instructional cues and the choice cues evoked a distributed network of regions in the frontoparietal cortex (Figure 2A). In particular, the dorsal attention network (DAN), comprising the intraparietal cortex/superior parietal lobule (IPS/SPL) and the frontal eye fields (FEF), was activated by both types of cues. To identify the brain networks preferentially engaged during willed attention control, we contrasted the choice cue-evoked BOLD activations with the instructional cue-evoked BOLD activations. This is shown in Figure 2B, where this contrast identified a network comprising the bilateral anterior insula (AI), bilateral anterior prefrontal cortex (APFC), bilateral dorsal lateral prefrontal cortex (DLPFC), dorsal anterior cingulate cortex (dACC), and bilateral inferior parietal lobule (IPL); the results were consistent across the two datasets. These brain regions, summarized in Table 2, were used to define ROIs for further analysis below. In addition, the dorsal attention network does not appear in the choice > instructed contrast maps, indicating that its cue-evoked activation did not differ detectably between externally instructed and internally willed attention under the present analysis. See (Rajan et al., 2019) and (Bengson et al., 2015) for similar analysis on these datasets.

**Figure 2.**
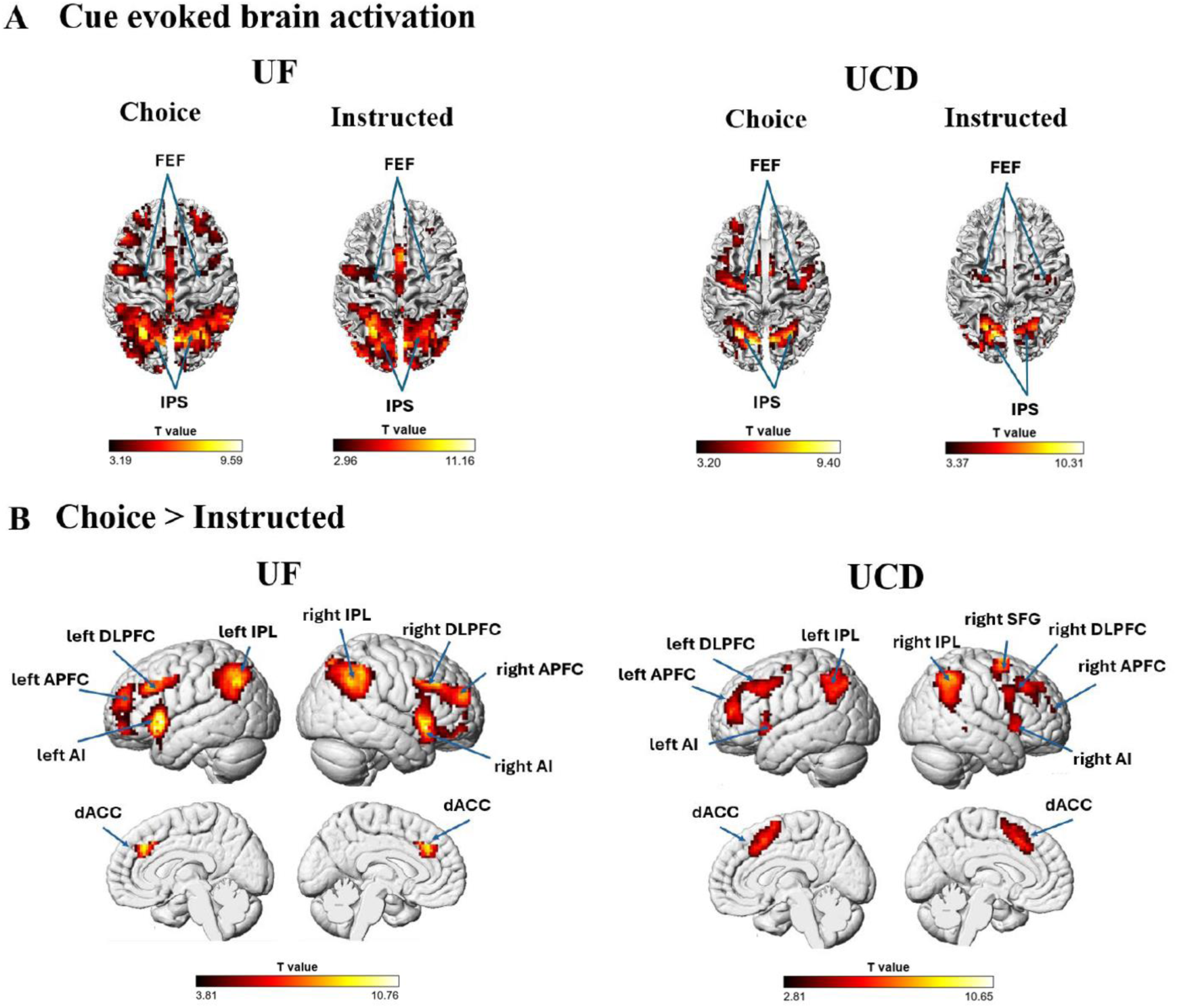
Univariate BOLD activation analysis. (A) The DAN was activated by both the instructional cues and the choice cue in both datasets. (B) Choice cue, however, additionally activated a frontoparietal network consisting of AI, APFC, DLPFC, dACC, and IPL in both datasets. (AI: anterior insula; APFC: anterior prefrontal cortex; DLPFC: dorsal lateral prefrontal cortex; dACC: dorsal anterior cingulate cortex; IPL: inferior parietal lobule; SFG: superior frontal gyrus). The statistical maps were thresholded at p < 0.05 FDR.

### Multivariate analysis of cue-evoked BOLD activation

To test whether neural activity patterns in the frontoparietal ROIs differentiated the two attention conditions, MVPA was applied to each ROI to decode attend-left versus attend-right for instructional trials and choice trials separately; see Figures 3A and 3B. For both datasets, decoding accuracy for the instructed trials was not significantly different from chance level (50%) for all the ROIs. For the choice trials, decoding accuracy was significantly above chance level only for a small number of ROIs in the individual datasets (1 for UF dataset) and (3 for UCD dataset). To increase statistical power, we combined the data from the UF dataset and the UCD dataset via meta-analysis, and the result is shown in Figure 3C. For the choice trials, decoding accuracy between attend-left and attend-right conditions was significantly above chance in all the ROIs at p ≤ 0.05 FDR, but for the instructed trials, it remained at chance level in all the ROIs. These results demonstrate that the network of frontal and parietal brain regions preferentially activated during willed attention is involved in deciding (choosing) the direction of attention (left vs. right). In contrast, during instructed attention, the same ROIs did not contain decodable information about the direction of attention. These decoding findings are further corroborated using a permutation-based statistical analysis (Combrisson & Jerbi, 2015). See Table A1, where effects sizes are also reported. We henceforth refer to the frontoparietal regions preferentially activated by the choice cue as the frontoparietal decision network.

**Figure 3.**
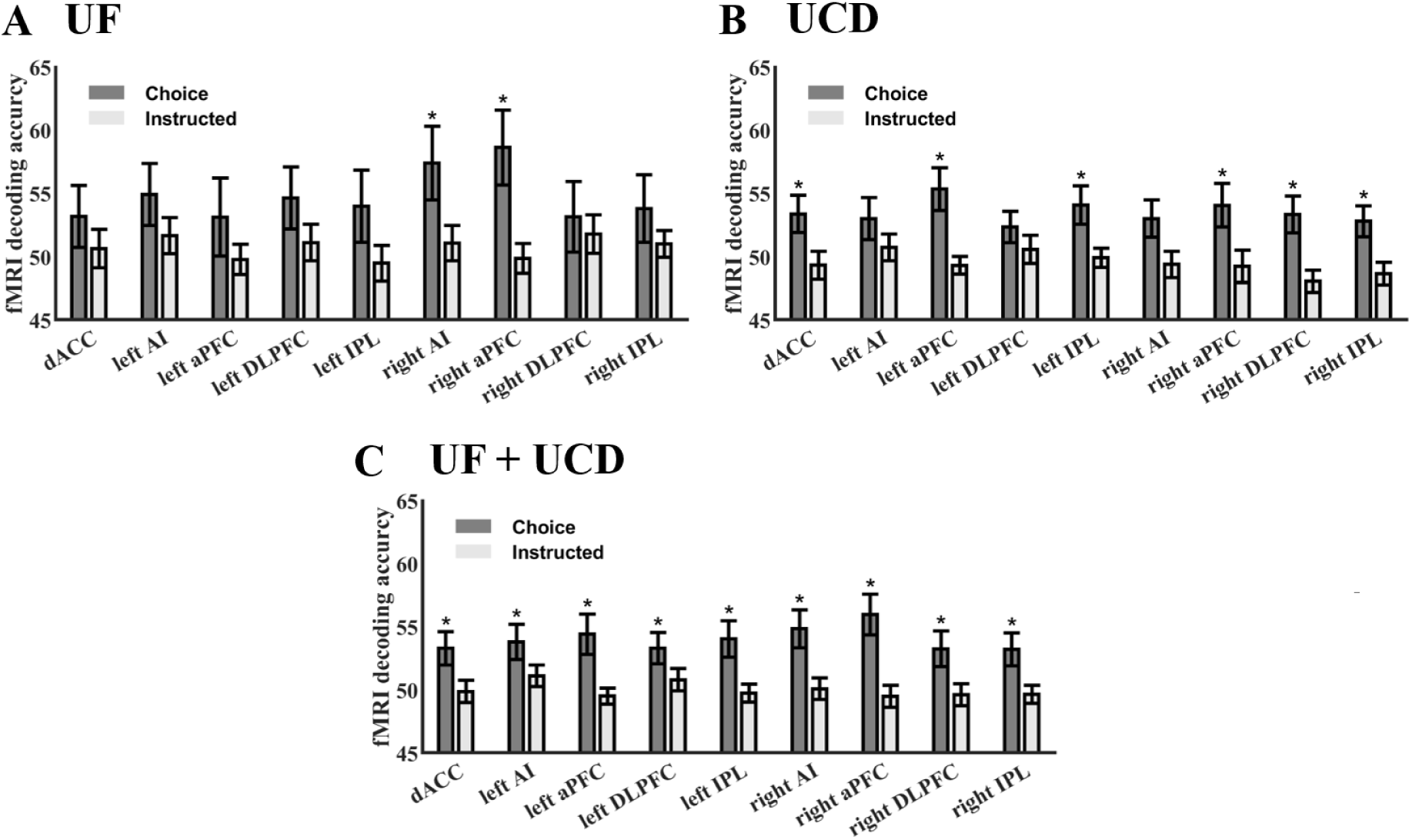
Decoding accuracy between attend-left and attend-right for instructed and choice trials in frontoparietal ROIs during the cue-to-target period. (A) UF dataset. (B) UCD dataset. (C) Combining UF and UCD via meta-analysis, it was observed that in all 9 ROIs, decisions about where to attend could be decoded in choice trials but not in instructed trials. *: p≤0.05 FDR.

### Precue alpha power predicts the decision about where to attend

For both the UF and UCD datasets, precue alpha power in the interval from –500 ms to 0 ms was extracted from each electrode and subjected to MVPA analysis to examine whether it predicted the postcue direction of attention (Figure 4A). In the UF dataset, decoding accuracy based on precue alpha power was 55.74% for the choice condition (p = 0.04), significantly above chance level. In contrast, decoding accuracy for the instructed condition was 49.57%, which did not exceed chance (p = 0.67). Similarly, in the UCD dataset, decoding accuracy was 57.29% for the choice condition (p < 0.0001), again significantly above chance, and 49.22% for the instructed condition (p = 0.46), which was not above chance. In Figure A1, it is further shown that this pattern of results holds if we only use the posterior electrodes or the frontal electrodes, suggesting that both posterior and frontal channels can contribute to decoding the upcoming decision about where to attend. Together, these highly consistent results across two independent datasets demonstrate that multichannel patterns of EEG alpha power in the –500 to 0 ms interval prior to the onset of the choice cue predicted the direction of attention during willed attention, whereas, as expected, alpha patterns preceding instructional cues did not. These findings from MVPA analysis support and further substantiate the findings of Bengson et al. (2014).

**Figure 4.**
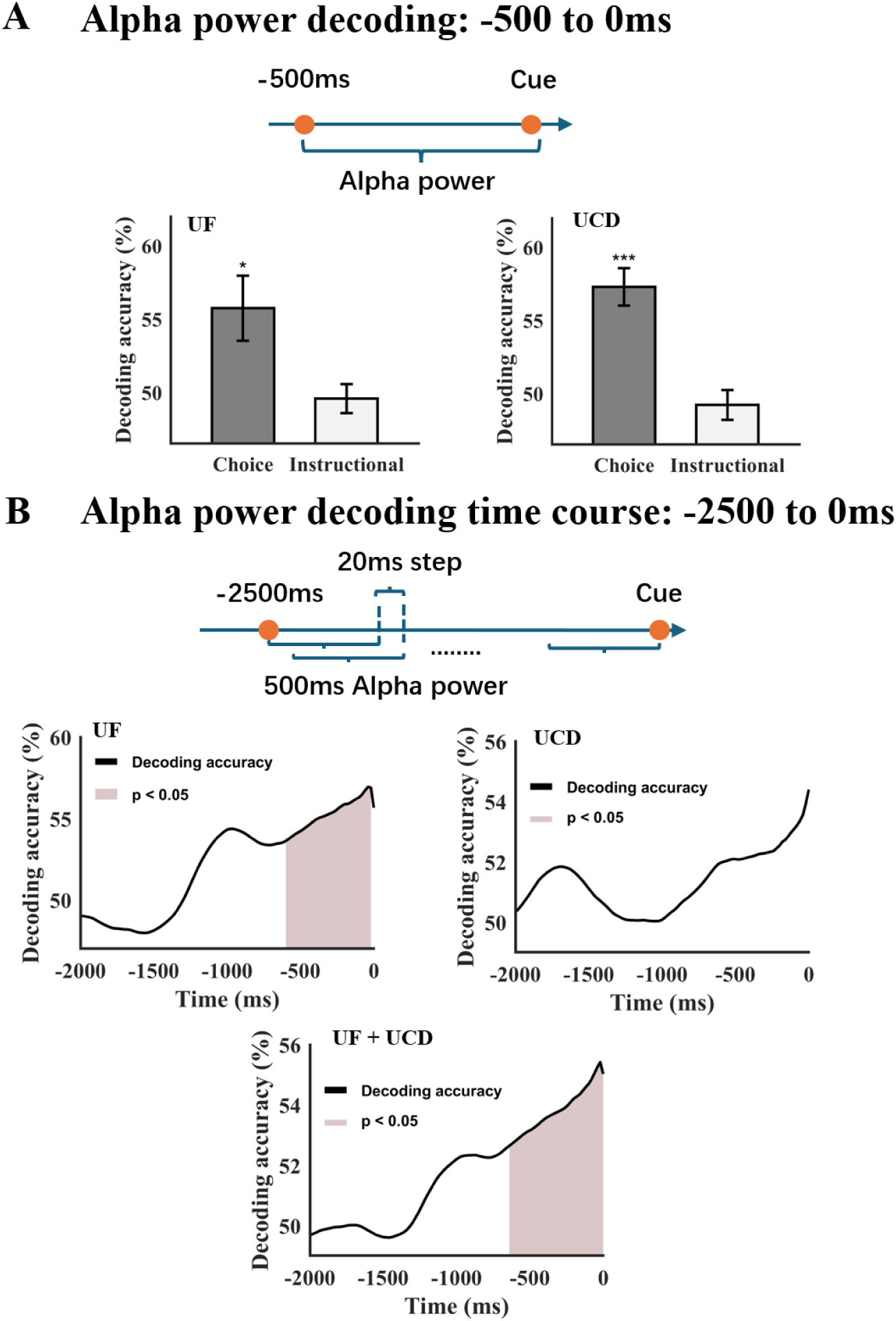
Precue alpha power patterns and postcue direction of attention. (A) Alpha power decoding accuracy (−500 to 0 ms) for UF and UCD datasets. For both datasets, the postcue direction of postcue attention can be decoded for choice trials, but not for instructional trials. (B) Precue decoding accuracy time course suggests that only alpha activity immediately preceding the cue onset predicted the post cue direction of attention. *: p<0.05, ***: p<0.001.

To investigate whether the −500 ms to 0 ms alpha power pattern that predicted decisions about where to attend following the choice cue might have begun earlier in time, we performed a moving-window decoding analysis over the time period of −2500 ms to 0 ms where the duration of the moving window was 500 ms and the step size was 20 ms; the time of the window was labeled by the time of the rightmost point (e.g., the time label for the window −2500 to −2000 ms was “-2000 ms”). Figure 4B shows the results. Similar patterns were seen for both the UF and UCD datasets. Combining the two datasets, we found that from −2000 ms to −1200 ms, the decoding accuracy hovered around 50%, and starting at around −1200 ms, the decoding accuracy began to rise and became above chance level around −650 ms prior to the onset of the cue. A cluster-based permutation approach (Maris & Oostenveld, 2007) further showed that the cluster from −650 ms to 0 ms was significantly above chance controlling for family-wise error at p = 0.008. These results show that predictive information in precue alpha patterns was confined to the approximately 650 ms immediately preceding cue onset, rather than extending across the earlier precue interval.

### Relation between precue alpha and postcue frontoparietal activity

To investigate the relation between precue alpha power pattern and postcue frontoparietal brain activity during willed attention, we divided the subjects into two groups (median split)—a high precue alpha decoding accuracy group and a low precue alpha decoding accuracy group. We then computed the univariate BOLD activations and multivariate decoding accuracies for each alpha power decoding accuracy group. For conciseness, in this analysis, we combined all the ROIs in Table 2 into a single frontoparietal decision ROI. Figure 5A shows the difference in BOLD activation between the two alpha power decoding-defined groups. For both the UF and UCD datasets, the subjects with high precue alpha decoding accuracy showed lower BOLD activation in the frontoparietal decision network. The difference was statistically significant in the combined UF/UCD dataset using meta-analysis. Figure 5B shows the difference in decoding accuracy between the two groups. In contrast to the pattern from the univariate BOLD activation results, the decoding accuracy in the frontoparietal decision network evoked by the choice cue was higher for participants with higher precue alpha decoding accuracy.

**Figure 5.**
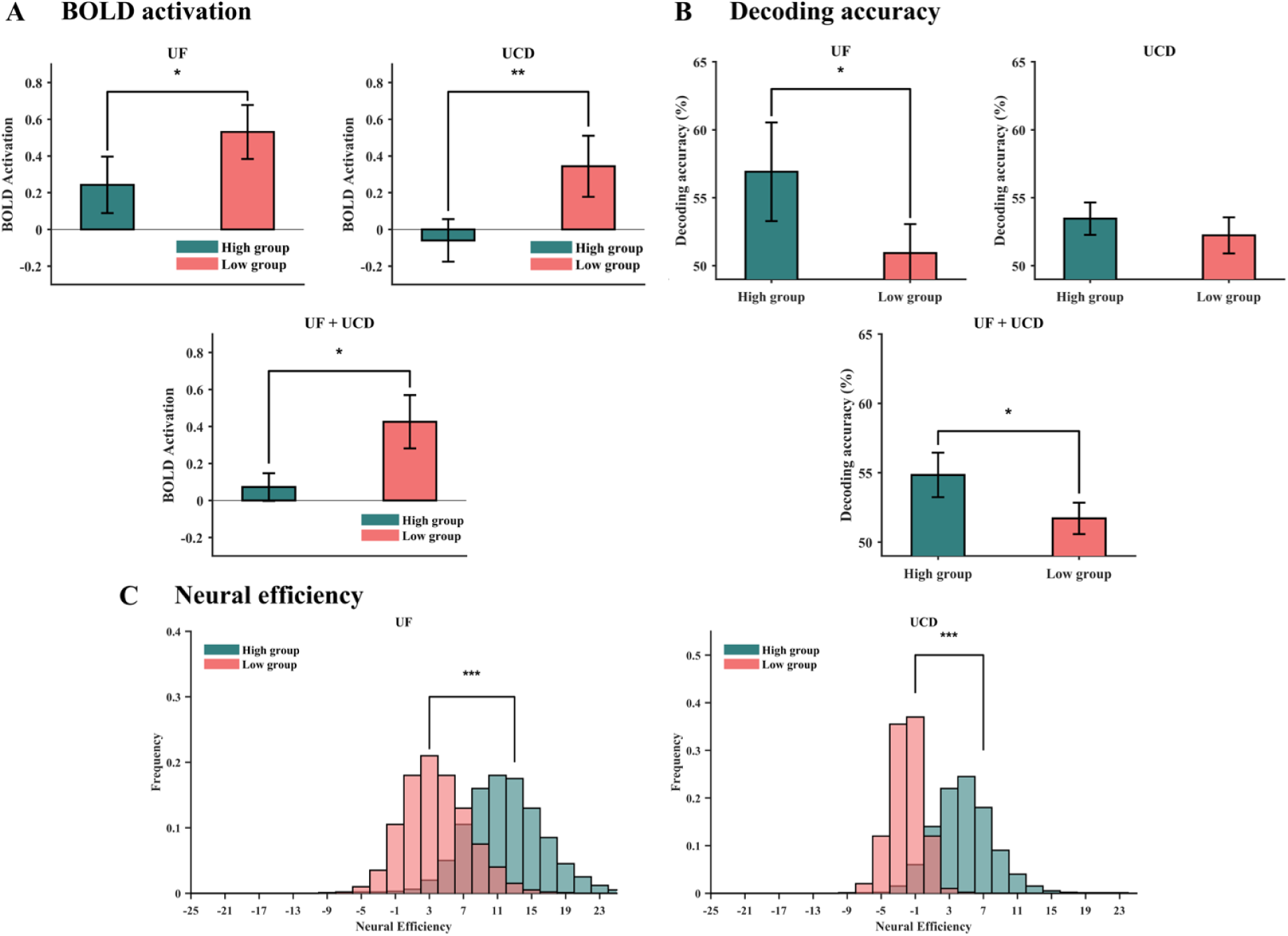
EEG informed fMRI analysis. Subjects were divided into two groups (median split) based on their precue alpha decoding accuracy being high or low. (A) The BOLD activation in the frontoparietal decision network for the two datasets (top) and meta-analysis combining the two datasets (bottom). (B) The decoding accuracy in the frontoparietal decision network for the two datasets (top) and meta-analysis combining the two datasets (bottom). (C) The bootstrap distributions of the neural efficiency for the two groups for the two datasets. * p<0.05, ** p<0.01, ** p<0.001.

Based on the results in Figures 5A and 5B, we computed an exploratory neural efficiency index in the frontoparietal decision network (Figure 5C), defined as the difference between the decoding accuracy for the choice trials and that for the instructed trials divided by the difference between the univariate BOLD activation for the choice trials and that for the instructed trials (see Methods). This definition is motivated by considering the decoding accuracy as an index of the strength of the decision about where to attend and the univariate BOLD activation as an index of the magnitude of the choice-related neural response; stronger decisions being reached at smaller neural responses are considered more neurally efficient. We applied bootstrap resampling to each of the two groups of subjects (high vs low precue alpha decoding accuracy) to obtain a distribution of 10,000 neural efficiencies. For the UF dataset, the mean neural efficiency = 12.18 for the high precue decoding accuracy group and the mean neural efficiency = 3.89 for the low precue decoding accuracy group; for the UCD dataset, the mean neural efficiency = 4.51 versus −2.02 for the high vs low precue decoding accuracy groups. The high vs low means were significantly different, as the two distributions differed significantly according to the Kolmogorov–Smirnov test (p < 0.0001), and the result was consistent across both datasets. This precue-postcue analysis indicates that participants with more predictive precue alpha patterns showed higher values of the neural efficiency index. One cautionary note is that, because ratio-based measures can be unstable when their denominator is small or noisy, this index should be treated as a suggestive measure rather than the main outcome measure. Additional research is needed to further establish its robustness and validity.

## Discussion

In this study, we analyzed fMRI and EEG data from two independent willed attention datasets, acquired using the same paradigm, to address the following questions. What is the neural network underlying willed attention control? What is the functional role of this willed attention control network? How does the neural activity prior to willed decisions about where to attend influence brain activations related to willed attention control? We found that (1) relative to instructed attention, during willed attention control, there were higher BOLD activations in regions of the frontal and parietal cortex, including in the dorsal anterior cingulate (dACC), bilateral anterior insular (AI), bilateral anterior prefrontal cortex (APFC), bilateral dorsolateral prefrontal cortex (DLPFC), and bilateral inferior parietal lobule (IPL), (2) choices to attend left versus attend right could be decoded from the neural activity patterns in these frontoparietal regions, but this was not true for instructions to attend left versus attend right, (3) multichannel patterns of EEG alpha power immediately prior to the onset of the choice cue predicted subsequent decisions about where to attend, but this was not true for instructed attention, and (4) subjects with higher precue alpha decoding accuracy had higher values of the neural efficiency index, defined as the ratio of the choice-versus-instructed difference in decoding accuracy to the corresponding difference in univariate BOLD activation..

### Role of the frontoparietal decision network

Willed attention is volitional behavior. As such, some parallels can be drawn with volitional movements (actions), particularly in the context of freely choosing among multiple motor alternatives. Studies of volitional movements have consistently revealed that the anterior cingulate cortex (ACC) serves as a critical component of the volitional control network, playing essential roles in voluntary response selection, conflict monitoring, and evaluation of outcomes of self-generated decisions (Rushworth et al., 2011; Walton et al., 2004). Similarly, the dorsolateral prefrontal cortex (DLPFC) has been identified as crucial for choosing between alternative predefined movements, particularly in tasks requiring selection among multiple alternatives (Frith et al., 1991; Jahanshahi et al., 1995). Remarkably, in willed attention, a cognitive act, a similar frontoparietal network is activated by the choice-cue processing (Bengson et al., 2014; Liu et al., 2017; Rajan et al., 2019). Although earlier work employing univariate analysis has reported similar findings (Greenberg et al., 2010; Hopfinger et al., 2010; Taylor et al., 2008), our contribution, employing multivariate pattern analysis (MVPA), substantially extends the prior work by directly demonstrating that the frontoparietal regions activated by the choice cue encode decision-relevant information, even in the absence of overt motor output. This led us to refer to these frontoparietal regions collectively as the frontoparietal decision network. One might be tempted to link the findings to the premotor theory of attention and view the differential activation patterns as reflecting potential movement plans. This, however, is unlikely, as the motor response requirement is the same for both instructed and willed attention and we observed no differential patterns between attend left and attend right in the instructed (cued) trials.

Network neuroscience has suggested that the frontoparietal decision network can be further divided into two complementary networks: the central executive network (CEN) and the salience network (SN). The CEN, anchored in the dorsolateral prefrontal cortex (DLPFC) and inferior parietal lobule (IPL), is central to deliberative processes such as weighing multiple alternatives, integrating task-relevant information, and executing context-appropriate choices (Niendam et al., 2012; Walsh et al., 2011). When faced with multiple movement options, the CEN employs competitive accumulation mechanisms where each potential choice has a corresponding neural accumulator that simultaneously accumulates evidence until one reaches the decision threshold (Cisek & Kalaska, 2010). The SN, comprising the anterior insula (AI) and dorsal anterior cingulate cortex (dACC), supports the identification of behaviorally salient cues and initiates cognitive shifts needed to implement decisions (Menon & Uddin, 2010; Seeley et al., 2007). In multi-choice volitional movements, the SN coordinates competitive inhibition between alternative motor programs, with increased lateral inhibition mechanisms required to manage competition among multiple options rather than simple binary choices (Cisek, 2007). Our results contribute to the growing literature by demonstrating that in willed attention, where overt motor-response requirements were matched across conditions, these two networks form an integrated decision-making architecture that mirrors multi-choice motor control. The anterior prefrontal cortex (aPFC), while it is not typically associated with the two networks, being situated at the apex of the prefrontal hierarchy, it is thought to maintain higher-order abstract goals to top-down bias decision-making processes (Badre & D’Esposito, 2009).

It is worth noting that the present results do not fully decompose the distinct functional contributions of the component regions of the frontoparietal decision network. Regions such as the dACC, AI, DLPFC, IPL, and aPFC participate in a broad range of processes, including cognitive control, working memory, task monitoring, salience processing, and responses to task difficulty. Thus, although the choice-related activation and left-versus-right decoding findings link this network to the internally generated selection of an attentional direction, they do not establish a unique role for any individual node or distinguish all of the processes that contribute to willed attention. The recruitment of AI and dACC during choice trials also raises the possibility that willed attention involves greater effort or arousal-related engagement than instructed attention. Consistent with this possibility, choice trials showed numerically slower RTs and higher accuracy than matched instructed trials, a pattern consistent with a speed-accuracy trade-off, although these behavioral differences were not consistently significant across measures and sites. In addition, postcue occipital alpha suppression was stronger for choice than instructed trials in the UF dataset, with a similar but nonsignificant pattern in the UCD dataset (see Figure A2). Because pupillometry was not recorded and alpha suppression is not a specific measure of arousal, these observations do not establish an arousal mechanism. They do, however, support the possibility that the additional frontoparietal recruitment during willed attention reflects increased engagement that helps maintain performance while participants generate their own attentional goal.

### Influence of precue brain state on postcue decision-making

Decision-making is under the influence of both external and internal factors. In willed attention, because the same choice cue prompts the two possible decisions (attend-left or attend-right), the external environment does not bias the choice being made. The internal environment defined by the momentary state of the brain, however, fluctuates from trial to trial. It is reasonable to hypothesize that such fluctuations prior to the onset of the choice cue might contribute to bias the decisions prompted by the choice cue. In a perceptual guessing task, in which the participants guessed the object category of the noisy input (Bode et al., 2012), it was shown that EEG activity immediately preceding the onset of pure noise patterns predicted the participant’s guessed object category, demonstrating the biasing influence of the brain state on perceptual decision-making when the stimulus input contains no discriminating information.

It is known that alpha oscillations are an important index of ongoing brain state (Jensen & Mazaheri, 2010; Klimesch et al., 2007). In the context of decision-making, alpha power in specific brain regions has been related to decision outcomes in paradigms in which the stimulus contains information favoring one decision over another (Jensen & Mazaheri, 2010; Klimesch et al., 2007; Sadaghiani & Kleinschmidt, 2016). Our approach differs from previous work in several aspects. First, our multivariate approach captures distributed spatial patterns across multiple electrodes simultaneously (Grootswagers et al., 2017; King & Dehaene, 2014), potentially revealing decision-relevant information that could be missed by examining single channels in isolation. Second, our paradigm represents a significant departure from conventional decision-making research (Britten et al., 1992; Gold & Shadlen, 2007; Newsome & Pare, 1988; Ratcliff & McKoon, 2008) by having the same choice cue prompt two different decisions, eliminating the influence of stimulus-driven evidence accumulation and placing more emphasis on the role of internal brain state fluctuations. Our findings demonstrate that multivariate alpha patterns predict upcoming choices in the absence of external evidence favoring either spatial alternative. Predictive alpha information was confined to the approximately 650 ms preceding cue onset rather than extending across the earlier precue interval. Complementary EEG evidence from the same paradigm showed that choice-specific frontal theta activity emerges approximately 400–800 ms after the choice cue and that posterior alpha lateralization, an index of attentional-set implementation, is delayed by approximately 400–500 ms in choice trials relative to instructed trials (Rajan et al., 2018). Together, these precue and postcue temporal dynamics indicate that the principal decision process occurs near and after presentation of the choice cue, rather than being maintained throughout the preceding intertrial interval.

These findings extend Bengson et al. (2014) in several important ways. Bengson et al. showed that lateralized parieto-occipital alpha power in the 1-s interval preceding the choice cue was related to the subsequent direction of willed attention. Here, using multivariate decoding of distributed alpha-power patterns, we show that the upcoming choice can be predicted from multichannel alpha activity in two independent datasets. The present moving-window analysis further refines the temporal profile of this effect: predictive information was concentrated in the approximately 650 ms immediately preceding cue onset rather than being distributed across the earlier precue interval. Finally, by relating individual differences in precue alpha decoding to postcue frontoparietal activation and decoding, the present study links ongoing alpha-state patterns to subsequent decision-related frontoparietal activity (see below).

### Linking precue activity to postcue response

In the foregoing, we discussed postcue frontoparietal encoding of decision-related variables and precue brain state biasing of postcue decision-making. How might the two relate to one another? We addressed this question by showing that individuals whose precue brain states more effectively predicted their postcue decisions demonstrated enhanced encoding of decision variables within the choice-activated frontoparietal decision network (i.e., higher decoding accuracy). To understand the possible neural bases of this effect, let us consider the role of the dACC, which is known to play an important role in monitoring the cognitive environment, detecting conflicts, and directing the central executive network to exert top-down modulation of sensorimotor processing (M. M. Botvinick et al., 2001; Carter et al., 1998; Shackman et al., 2011). Whereas early work emphasized the dACC’s role in the monitoring of the external environment (Bush et al., 2000; Devinsky et al., 1995), later studies have begun to investigate the dACC’s role in monitoring the internal environment as well (Alexander & Brown, 2011; Heilbronner & Hayden, 2016; Shenhav et al., 2013). In particular, evidence suggests that dACC, as part of the broader frontoparietal decision network, monitors ongoing neural oscillatory patterns, including alpha activity (Jensen & Mazaheri, 2010; Klimesch, 2012; Sauseng et al., 2011). In addition, deciding between two equal alternatives represents a conflict situation, and the role of the dACC in conflict resolution is well-established (M. Botvinick et al., 1999; Kerns et al., 2004; Sheth et al., 2012). These considerations make it plausible that during choice trials, dACC monitors the internal brain state, indexed by the pattern of alpha oscillations, and resolves the conflict of choosing where to attend among equal alternatives under the influence of such monitoring. Importantly, our findings further extend the role traditionally assigned to dACC to the entire frontoparietal decision network (Cole & Schneider, 2007; Dosenbach et al., 2007; Vincent et al., 2008), consistent with the reports showing that frontoparietal areas, including dLPFC and posterior parietal cortex, contribute to monitoring uncertainty and implementing control adjustments based on assessment of internal state (Brass et al., 2005; Corbetta & Shulman, 2002; Hampshire et al., 2010). In addition to enhanced representation of the decision variables, participants whose precue alpha better predicted the postcue decision had lower BOLD activity to the choice cue in the frontoparietal decision network, suggesting that stronger postcue decisions may be reached at lower neural energy costs (Attwell & Iadecola, 2002; Neubauer & Fink, 2009). Introducing a new variable called neural efficiency to quantify this, we found that participants who can better incorporate the precue brain state into postcue decision-making have higher values of the neural efficiency index.

### A model of willed attention control

Both instructional and choice cues activated the dorsal attention network (DAN). When choice-cue-evoked BOLD signals were contrasted with those evoked by instructional cues, the DAN did not appear in the resulting brain map, indicating that cue-evoked DAN activation did not differ detectably between willed and instructed attention. In addition, in a recent study (Meyyappan et al., 2025), we showed that the direction of attention during the cue-target interval could be decoded in the DAN and throughout the retinotopic visual cortical hierarchy, irrespective of whether the covert attentional state was generated by external instructions or internal decisions. Together, these results suggest that willed attention control is achieved through the collaborative engagement of the frontoparietal decision network and the DAN. A proposed model is shown in Figure 6: (1) following the choice cue, the decision about where to attend is formed in the frontoparietal decision network under the influence of ongoing brain state; (2) this decision is transmitted to the DAN to initiate attention control; (3) the DAN issues top-down biasing signals to retinotopic visual cortex in anticipation of the visual stimulus; and (4) the neural mechanism in Step 3 is shared between willed and instructed attention.

**Figure 6.**
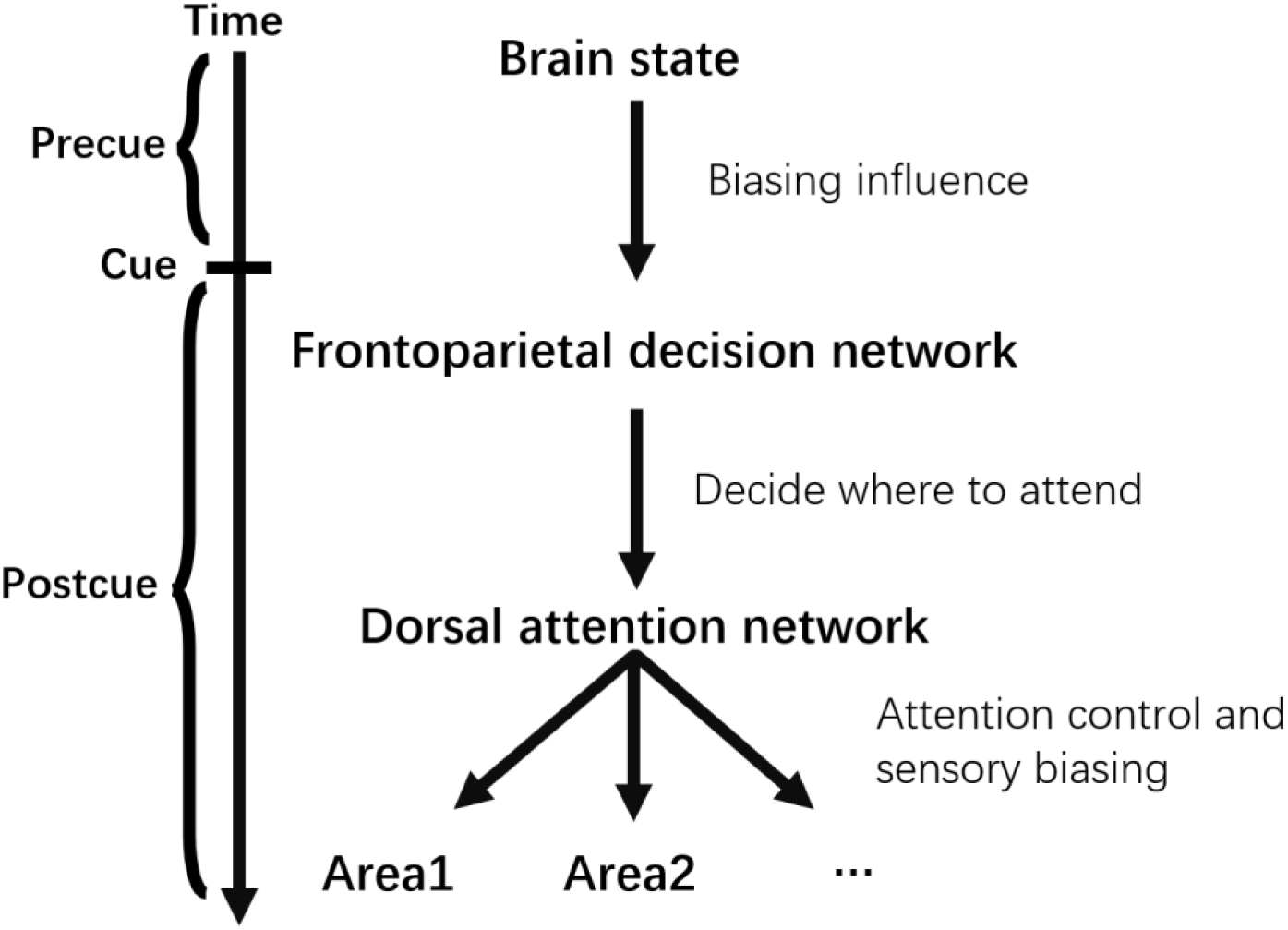
The model of willed attention control. The frontoparietal decision network, under the biasing influence of the ongoing precue brain state, makes the decision about where to attend upon reception of the choice cue and sends the decision to the dorsal attention network. The dorsal attention network executes attention control and issues top-down signals to bias visual areas in anticipation of the upcoming visual processing. Attention selection of sensory input occurs as the result of top-down biasing. Sensory biasing and stimulus section mechanisms are shared between willed and instructed attention.

### Limitations

Several limitations should be considered. First, the willed-attention paradigm captures a constrained instantiation of volitional attention: participants chose between two spatial locations after presentation of an explicit choice cue. More natural forms of willed attention, including self-initiated shifts without an external prompt, decisions about which object or feature to attend, decisions about when to initiate or terminate an attentional episode, and attention guided by personally meaningful goals, remain to be explored (Nadra & Mangun, 2023). Future studies should examine whether the frontoparietal decision network and precue alpha effects identified here generalize to these broader forms of volitional attention. Second, because choice trials were classified according to participants’ self-selected attend-left and attend-right responses, individual participants could show unequal numbers of trials in the two classes. We examined this issue in Figure A3. After equating the numbers of choose-left and choose-right trials within each participant and evaluating performance against an empirical within-subject label-permutation distribution, decoding remained significantly above the permutation baseline. Thus, the principal precue-alpha result was not explained by trial imbalance or subject-specific left-right choice biases. Nonetheless, future work should use designs that yield larger numbers of trials and directly model individual choice biases and sequential strategies. Third, the neural efficiency index, as a ratio measure, can be sensitive to small or noisy denominators. The index should therefore be interpreted as a suggestive descriptive measure rather than a standalone biomarker of decision efficiency. Future work should consider complementary multivariate approaches that model decoding and activation separately rather than combining them as a ratio, while accounting for other task- and participant-level variables that may influence either measure.

## Acknowledgements

This work was supported by NIH grant MH117991 and NSF grant BCS2318984.

